# Plant responses to butterfly oviposition partly explain preference-performance relationships on different brassicaceous species

**DOI:** 10.1101/706044

**Authors:** Eddie Griese, Ana Pineda, Foteini G. Pashalidou, Eleonora Pizarro Iradi, Monika Hilker, Marcel Dicke, Nina E. Fatouros

## Abstract

According to the preference-performance hypothesis (PPH), also known as ‘mother-knows-best hypothesis’, herbivorous insects prefer those plants for oviposition, which yield the best offspring performance. Yet, most studies testing the PPH neglect the possibility that plant responses to insect eggs may affect both egg survival and larval performance. Here, we tested the PPH by studying responses of seven Brassicaceae plant species to oviposition by two cabbage white species. When including the egg phase, our study supports the ‘mother-knows-best hypothesis’: larvae of *Pieris rapae* (solitary) or *P. brassicae* (gregarious) gained most weight on those plant species which had received most eggs (*B. nigra* or *B. montana*, respectively). However, our experiments did not reveal any relationship between oviposition preference and egg survival. Brassicaceous species are known to respond to these butterfly eggs with a hypersensitive response (HR)-like necrosis, which can lower egg survival. *Pieris* eggs frequently induced necrosis in five of the tested plant species. Survival of clustered *P. brassicae* eggs was unaffected by HR-like in four of the five species. Therefore, our experiments did not reveal any relationship between *P. brassicae* egg survival and oviposition preference. Females of *P. rapae* preferred oviposition on plant species which most frequently showed HR-like necrosis. Remarkably, although egg survival was lower on HR-like plants, larval biomass was higher compared to plants without a necrosis. We conclude that egg survival does not seem to be a deciding factor for oviposition choices. However, egg-mediated plant responses might be important to explain the PPH of the two *Pieris* species.

**Lay summary:** Egg-laying preferences of herbivorous insects can often be linked to offspring performance. Commonly, the fate of insect eggs and the plant responses to the eggs have been ignored when studying the link between preference and performance. By including the egg phase, our study supports the ‘mother-knows-best hypothesis’, explained by butterfly oviposition and associated egg and larval performances on different plant species. We especially found that egg-mediated responses seem a deciding factor for butterfly oviposition choices.

## Introduction

Host-plant selection for oviposition by insect females is a decisive step in establishing a new herbivore generation (Gripenberg et al., 2010; Thompson, 1988a, b; Thompson and Pellmyr, 1991). The preference-performance hypothesis (PPH) or ‘mother-knows-best’ hypothesis states that natural selection favors those insect females which prefer host plants where the offspring performs best, especially when immature stages are less mobile than adults. A good host plant is usually characterized either by high food quality and/or by enemy-free space (Craig and Ohgushi, 2002; Gripenberg et al., 2010; Jaenike, 1990; Mayhew, 1997, 2001). Indeed, the PPH is supported by numerous studies of butterflies and moths (Forister, 2004; Forister et al., 2009; Harris et al., 2001; Thompson, 1988a, b; Thompson and Pellmyr, 1991). However, there are also numerous studies of plant - Lepidoptera interactions, where no support was found for the PPH (Gripenberg et al., 2010; Jaenike, 1990; König et al., 2016; Mayhew, 1997; Scheirs et al., 2000; Thompson, 1988a). In addition to host plant quality and presence of natural enemies, various factors such as local host plant abundance or distribution patterns of host plants shape oviposition preferences and larval performance (Friberg et al., 2015; Wiklund and Friberg, 2008, 2009).

Yet, the vast majority of innumerable laboratory studies testing the PPH did not consider that plants can activate defenses in response to egg deposition. Research has provided evidence that numerous plant species across highly diverse taxa defend against egg depositions of various insect species (Hilker and Fatouros, 2015). Plants are capable of killing eggs (Fatouros et al., 2016). For example, egg-induced formation of a neoplasm (Petzold-Maxwell et al., 2011) or by hypersensitive response (HR)-like necrosis at the oviposition site (Fatouros et al., 2014; Griese et al., 2017; Shapiro and DeVay, 1987) may result in egg detachment from the plant or egg dehydration. Furthermore, a plant can kill insect eggs by growing tissue that is crushing the eggs (Aluja et al., 2004; Desurmont and Weston, 2011; Karban, 1983; Mazanec, 1985). In addition, plants receive insect eggs as early ‘warning cues’ of impending herbivory and reinforce or prime their defenses against the subsequently feeding larvae (Austel et al., 2016; Bandoly et al., 2015; Hilker and Fatouros, 2015, 2016; Hilker et al., 2016; Pashalidou et al., 2015c; Pashalidou et al., 2013). As a consequence, larval performance on initially egg-infested plants may be worse than on egg-free plants (Hilker and Fatouros, 2016). Additionally, but less frequently shown so far, egg deposition can suppress plant defenses against larvae (Reymond, 2013). The effects of egg deposition on subsequent plant defenses against larvae that hatch from the eggs have been extensively overlooked until recently, with most studies on larval performance being conducted by placing larvae onto an egg-free host plant (Hilker and Fatouros, 2015, 2016).

The insect oviposition mode can have a significant impact on egg survival and larval performance. When eggs are laid in clusters, neonate larvae often show gregarious feeding behavior, which benefits offspring performance in some insect species (Allen, 2010; Clark and Faeth, 1997; Clark and Faeth, 1998; Denno and Benrey, 1997; Desurmont and Weston, 2011; Desurmont et al., 2014; Fordyce, 2003; Martínez et al., 2017; Wertheim et al., 2005). On the other hand, many herbivorous insects lay single eggs, spreading them over a larger area, possibly as a means of reducing predation risk and competition (Nufio and Papaj, 2001; Root and Kareiva, 1984). Egg-induced plant defense affecting larval performance is especially known for insect species laying eggs in clutches (Hilker and Fatouros, 2015). However, also the plant’s response to singly laid eggs of *Manduca sexta* reinforces the defense against *M. sexta* larvae (Bandoly et al., 2016). It remains to be elucidated whether the egg laying mode (single eggs vs. egg clutches) affects egg-induced plant defense targeting the eggs and how this in turn depends on the plant species receiving the eggs.

The aim of this study is to elucidate whether the PPH still holds when not only considering relationships between oviposition preference and larval performance, but also when including egg survival rates and egg-induced changes in plant suitability for feeding caterpillars. Therefore, we investigated oviposition preference, egg survival and larval performance of *Pieris brassicae* and *P. rapae* on eight Brassicaceae species. Pierid butterflies have co-evolved since 90 million years ago with their host plants in the order Brassicales (Edger et al., 2015; Wheat et al., 2007). Both butterfly species are known to use various wild and cultivated brassicaceous plants as hosts (Chew and Renwick, 1995; Feltwell, 1982; Gols et al., 2011), whereby *P. rapae* can include also non-brassicaceous plants in their diet (Friberg et al., 2015). *Pieris* caterpillars are well adapted to Brassicaceae by their ability to detoxify glucosinolates, plant secondary metabolites characteristic for this plant taxon (Hopkins et al., 2009). While *P. rapae* lays single eggs on plants, *P. brassicae* lays egg clutches containing up to 200 eggs (Feltwell, 1982). Egg deposition by these pierid species is known to induce an HR-like leaf necrosis in several host plant species (Fatouros et al., 2016). Additionally, previous egg deposition by *P. brassicae* on brassicaceous plant species was shown to negatively affect larval performance (Bonnet et al., 2017; Geiselhardt et al., 2013; Lortzing et al., 2018; Pashalidou et al., 2015a; Pashalidou et al., 2015b; Pashalidou et al., 2015c; Pashalidou et al., 2013). It remains unknown so far how egg deposition of the conspecific solitary species *P. rapae* affects subsequently feeding larvae through egg-mediated plant responses.

We specifically addressed the following questions: (1) Do females of the two pierid species prefer to oviposit on plants on which their eggs show highest survival rates and larvae perform best? (2) Is this oviposition choice affected by the plant species’ capability to activate an egg-killing response (i.e. HR-like necrosis)? (3) Is the butterflies’ oviposition choice affected by plant responses to oviposition that subsequently affect feeding larvae, e.g. egg-mediated priming of defenses? (4) Does the egg-laying mode of the two pierid species affect their oviposition choice?

## Material and Methods

### Insects and plants

*Pieris brassicae* L. and *P. rapae* L. (Lepidoptera: Pieridae) were reared in a greenhouse compartment (21±1°C, 50 – 70% RH, L16:D8) on *B. oleracea* var. *gemmifera* L. plants. Female butterflies mated two to three days after eclosion. Their oviposition preferences were tested two days after mating. The females have a high egg load at this age and mating status (David and Gardiner 1962).

Eight different brassicaceous species were used in a preference experiment, seven in a performance experiment. Apart from *Raphanus sativus* L., all plant species were non-domesticated species. We obtained *R. sativus* from De Bolster seed company (The Netherlands), *Hirschfeldia incana* L. Lagr.-Foss. from the U.S., California, *Brassica nigra* L. from the Centre of Genetic Resources (CGN, Wageningen, the Netherlands) from an early flowering accession (CGN06619), *Sinapis arvensis* L. from Vlieland, in the north of The Netherlands, *B. montana* Pourr. from CGN (CGN18472 accession from Italy), *B. rapa* L. from Binnenveld, west of Wageningen (The Netherlands) and *B. oleracea* L. ‘Kimmeridge’ from the south coast of England. *Arabidopsis thaliana* (Col-0) was used only for the preference tests. Because of its small size, it was excluded from performance studies, as more than just the focal egg-induced plant would be needed to feed the caterpillars. All plants were in the non-flowering stage when tested, except *A. thaliana*, which already flowered. All plants were cultivated in pots filled with potting soil; they grew in a climate room (18 ± 4 °C, 60 – 80% RH, L16:D8). To use plants of similar biomass in the bioassays, *B. oleracea* was four weeks old, and all other plants were three weeks old at the time of infestation.

### Butterfly oviposition preference

To determine which plant species is preferred for oviposition, we simultaneously offered the above-mentioned eight plant species to a mated female butterfly. One individual of each plant species was placed into a mesh cage (75 x 75 x 115 cm). The plants were set up in a circle with the leaves not touching each other. The design was a randomized block with 18 replicates per butterfly species. The two butterfly species were tested in separate cages at different time points in a greenhouse compartment (23 ± 5 °C, 50-70% RH, L16:D8). After placing the plants inside a cage, one mated female butterfly was released. The number of *P. rapae* eggs or *P. brassicae* egg clutches, respectively, was counted on each plant three hours after release of the butterfly. Preliminary experiments showed that most butterfly females will make an oviposition choice within this time period.

### Plant treatments for performance tests

To assess egg survival rates and performance of larvae on previously egg-deposited plants, a plant individual of each species was infested with either *P. rapae* or *P. brassicae* eggs. Each plant was placed in a cage, which was located in a climate room (21 ± 1°C, 50-70% RH, L16:D8). The first fully developed leaf of each plant (fourth or fifth from the top) was exposed to either *P. brassicae* or *P. rapae* butterflies for egg deposition, while the rest of the plant was covered with a fine mesh. We limited the number of eggs deposited onto a plant to 20 eggs of *P. brassicae* (laid in a clutch) and to eight single *P. rapae* eggs per plant. Limiting egg deposition was done by observing the butterflies after introduction into the cages and removing them as soon as they had deposited the mentioned number of eggs. Those numbers were chosen to mimic naturally occurring egg numbers per plant (Fatouros et al., 2014; Feltwell, 1982). Occasionally extra laid eggs were immediately removed using a fine brush (see Pashalidou et al. (2013) for details). In total, seven to nine plant individuals per species were infested with *P. brassicae* eggs, and six to seven plant individuals per species received *P. rapae* eggs.

### Plant response to egg deposition, egg mortality and larval performance

To determine egg survival, we counted the number of larvae hatching from the twenty (*P. brassicae*) or eight (*P. rapae*) eggs deposited on a plant. To assess larval performance and the impact of the plant’s response to previous egg deposition on larval performance, we divided the neonate larvae hatching from egg-deposited plants into two groups. Half of them were placed back onto the previously egg-infested plant (labeled ‘egg and feeding’, EF) (on the adaxial side of the leaf where they hatched), and the other half was transferred to an egg-free plant (labeled ‘feeding’, F) plant of the same species and placed onto the adaxial side of the leaf as well. Three and seven days after hatching, caterpillar weight was measured on a microbalance (accuracy = 1 µg; Sartorius AG, Göttingen, Germany). We weighed each caterpillar individually, and afterwards the caterpillars were transferred back to their original position, on EF or F plants. Every EF and F plant was considered one replicate.

### Statistical analysis

Data on *P. rapae* oviposition preferences for host plants were analyzed by a generalized linear model (GLM) (poisson family), with plant species as fixed factor and the number of eggs per plant as response variable. The post-hoc analysis was performed using a linear hypothesis test (multcomp package). Because *P. brassicae* laid most eggs in a single clutch each time, only oviposition ‘yes’ or ‘no’ was scored when determining oviposition preferences. These data were analyzed by a GLM (binomial family) with the plant species as fixed factor, and the presence/absence of oviposition as response variable (post hoc test: linear hypothesis test).

Data on egg survival of each butterfly species were analyzed by a generalized linear mixed effect model (GLMM, lme4 package) with binomial distribution. The model included egg survival as response variable, and plant species, presence/absence of HR as well as the interaction between both variables were used as fixed factors. Date of infestation was used as random factor. A post-hoc analysis was conducted using linear hypothesis tests for plant species and interaction terms.

To evaluate whether oviposition preferences of a plant species can be linked to egg survival, we ran a correlation analysis by using Spearman correlation as well as linear regression to generate regression lines. We conducted this analysis first by relating the fraction of eggs (or egg clutches) laid on each plant species with the fraction of eggs surviving on each plant species. To elucidate the relationship between the plant’s ability to express HR-like necrosis and egg survival, we correlated the fraction of eggs laid on each plant species to the fraction of eggs surviving on those plants, which expressed HR-like necrosis in response to the eggs. To elucidate the relationship between the plant’s ability to express HR-like necrosis and oviposition preference, we linked the fraction of eggs or egg clusters laid to the fraction of plants expressing HR.

Data on caterpillar weight obtained on all plant species (subjected to prior egg deposition or not) were analyzed by using linear mixed effect models (LMM). We calculated the average caterpillar weight per plant. The logarithm of the mean caterpillar weight three or seven days after hatching was used as dependent variable, plant species and egg infestation as well as the interaction between them were used as independent variables, the random factor was the date of egg infestation. A post-hoc analysis was conducted using linear hypothesis tests for plant species and interaction terms. The effect of HR-like necrosis on caterpillar weight was tested by using the subset of plants infested with eggs and performing LMM on the logarithmic data of the mean caterpillar weight. Expression of HR-like necrosis, plant species as well as the interaction between both factors were included into the model. A post-hoc analysis was conducted by using linear hypothesis tests for plant species and interaction terms.

To detect possible links between butterfly oviposition preference and performance of three or seven-day-old caterpillars, a linear regression analysis was conducted. We conducted this analysis first by relating the fraction of eggs laid on each plant species with the weight of caterpillars on each plant species. Furthermore, we ran an analysis by relating the fraction of eggs laid on each plant species to the weight of caterpillars on (i) plants which received eggs prior to larval feeding (EF) and (ii) plants which were left without any eggs (F); thus, we aimed to test the hypothesis that females prefer to oviposit on plants with most modest (for *P. rapae:* putative) egg-mediated reinforcement of defense against the larvae. In addition, we analyzed the relationship between caterpillar weight and the fraction of eggs laid on plants expressing HR-like necrosis; thus, we aimed to test the hypothesis that females prefer to oviposit on plants whose HR-like necrosis has the most modest effect on the performance of their offspring. Finally, we tested the relationship between the fraction of plants expressing HR-like necrosis and caterpillar weight. Thus, we aimed to gain insight in whether the frequency of HR-like necrosis in response to the eggs relates to caterpillar weight.

All analyses were performed using R 3.3.2 (R Core Team, 2016).

## Results

### Oviposition preference

#### Gregarious species

Even though, there is a marginal effect of egg distribution for *P. brassicae* among the plants (χ^2^ = 19.65, df = 7, *P* = 0.04, GLM, Figure 2A), the post-hoc test did not reveal any significant differences (Supplementary Table 1). *Arabidopsis thaliana*, which was in the flowering stage (in contrast to all other plant species), did not receive any egg clutch by *P. brassicae* in this setup. Oviposition choices of *P. brassicae* were not correlated to plant fresh weight (S = 1720.7, ρ = 0.25, *P* = 0.24, Spearman correlation, Supplementary Figure S1).

**Figure 1:**
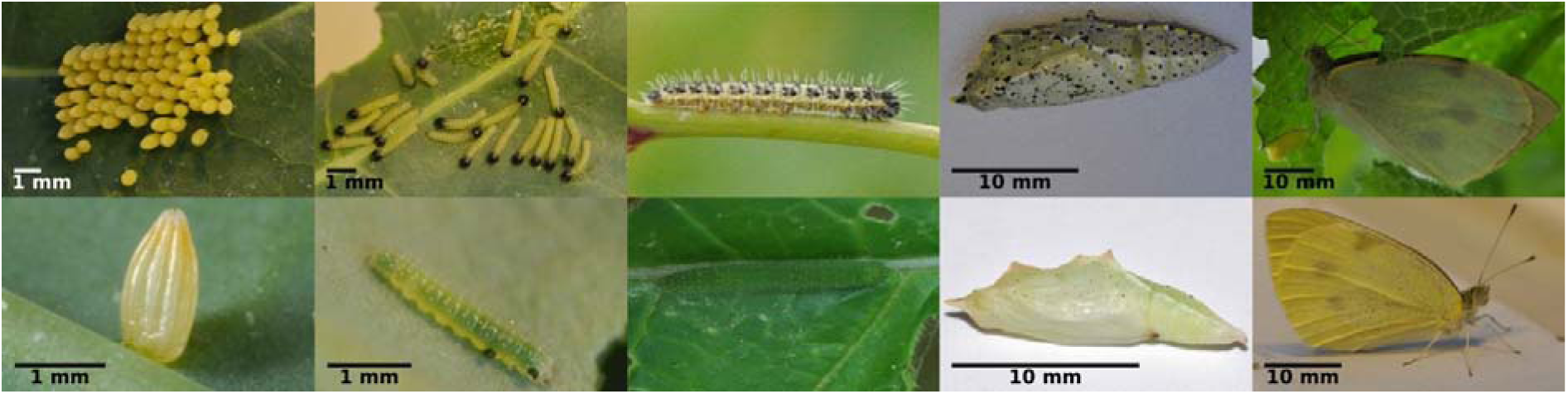
Developmental stages of the studied *Pieris* species. Differences in appearance and life history of the gregarious *P. brassicae* (top) and solitary *P. rapae* (bottom). From left to right: Eggs, neonates, L5 caterpillars, pupae, adults. *Pieris brassicae* larvae are feeding gregariously until the third larval stage. While eggs and adults of both species look similar and mainly differ in size, caterpillars of the two species are differently colored. Larvae of *P. brassicae* larvae are aposematically colored (an indication for unpalatability), which allows them to feed in groups, whereas *P. rapae* larvae are cryptically colored, a trait which is often linked with a solitary feeding behavior in butterflies (Sillén-Tullberg, 1988). Photo credits: E. Griese.

**Figure 2:**
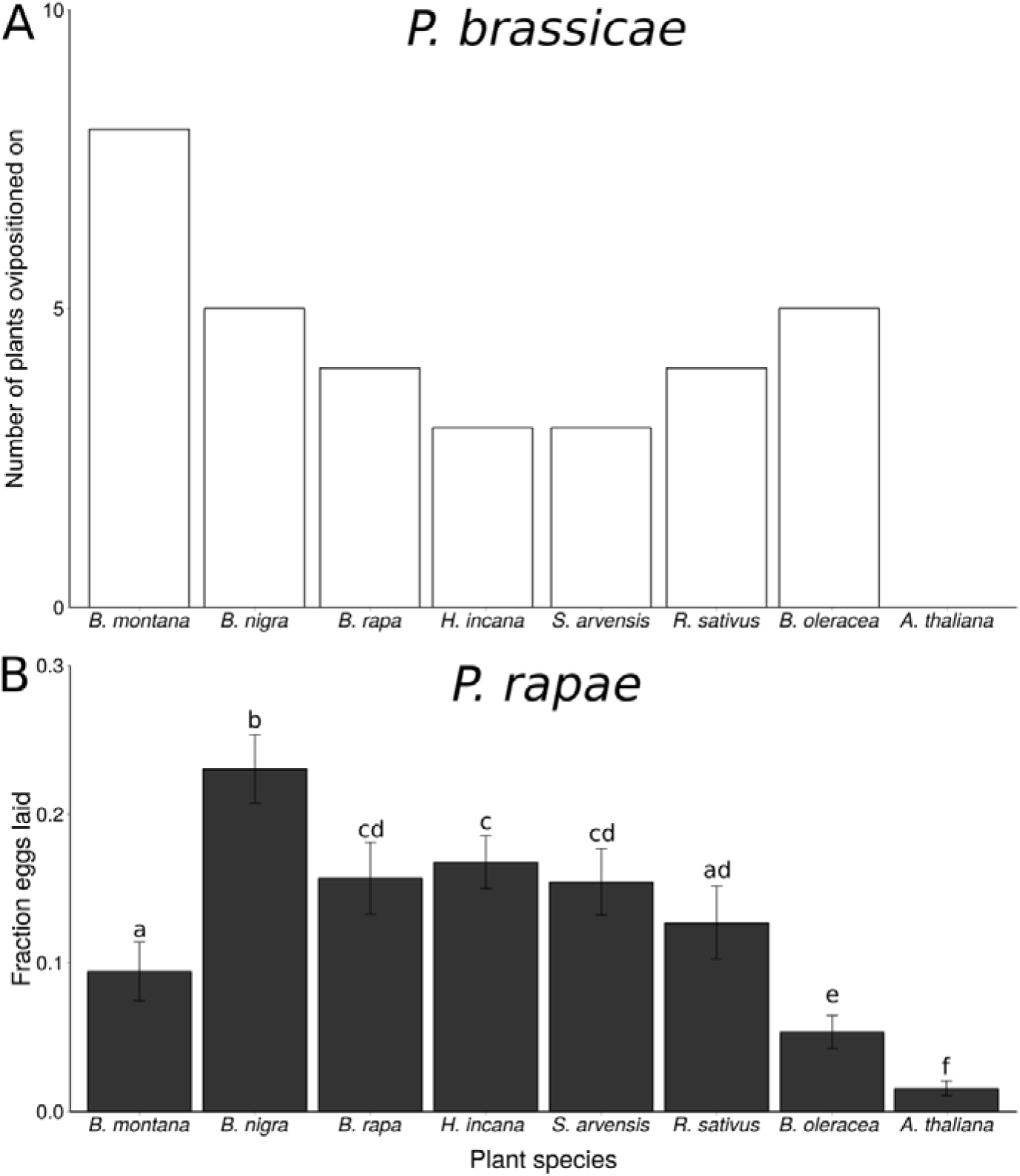
Oviposition preference bioassay. Fraction of plants, which received an egg deposition by *Pieris brassicae* (A) and fraction of eggs laid by *P. rapae* on different brassicaceous plant species (B). Mean fraction ± SE is given. (A) Female *P. brassicae* always laid at maximum of one egg clutch per plant; therefore, here the number of egg-deposited plants is given, whereas in (B) the number of single eggs of *P. rapae* were counted per plant species. In total, 18 plants per species were tested in random setups. Small letters indicate significant differences between plant species with *P* <0.05, GLM.

#### Solitary species

*Pieris rapae* females significantly preferred to oviposit on *B. nigra* over all other seven simultaneously offered plant species (Supplementary Table 1). The plant species chosen least frequently for oviposition were *B. oleracea* and *A. thaliana* (χ^2^ = 292.67, df = 7, *P* = <0.001, GLM, Figure 2B). The oviposition preference of *P. rapae* was not correlated to plant fresh weight (S = 1897.2, ρ = 0.18, *P* = 0.41, Spearman correlation, Supplementary Figure S1).

### Egg survival and effect of plant species and HR-like necrosis

#### Gregarious species

Egg survival was significantly affected by the plant species chosen by *P. brassicae* for egg deposition (χ^2^ = 20.39, df = 6, *P* = 0.002, GLMM, Figure 3A). Five out of the seven tested plant species expressed an HR-like necrosis in response to *P. brassicae* eggs (Figure 3A, Table 1). We observed the highest egg survival rates (almost 100%) when deposited on the plant species that were chosen most for oviposition (*B. montana, B. nigra).* Egg survival was significantly lower on *H. incana* and *R. sativus* compared to all other plants (apart from *B. nigra*) (Supplementary Table S2). Overall, induction of HR-like necrosis did not significantly affect egg survival (χ^2^ = 0.41, df = 1, *P* = 0.52, GLMM), while the interaction between the factors ‘HR’ and ‘plant species’ significantly affected survival of *P. brassicae* eggs (χ^2^ = 30.83, df = 4, *P* < 0.001, GLMM). On *B. montana*, egg survival was much lower on the two plants expressing HR than on the seven non-HR-plants (Figure 3A, Supplementary Table S3).

**Figure 3:**
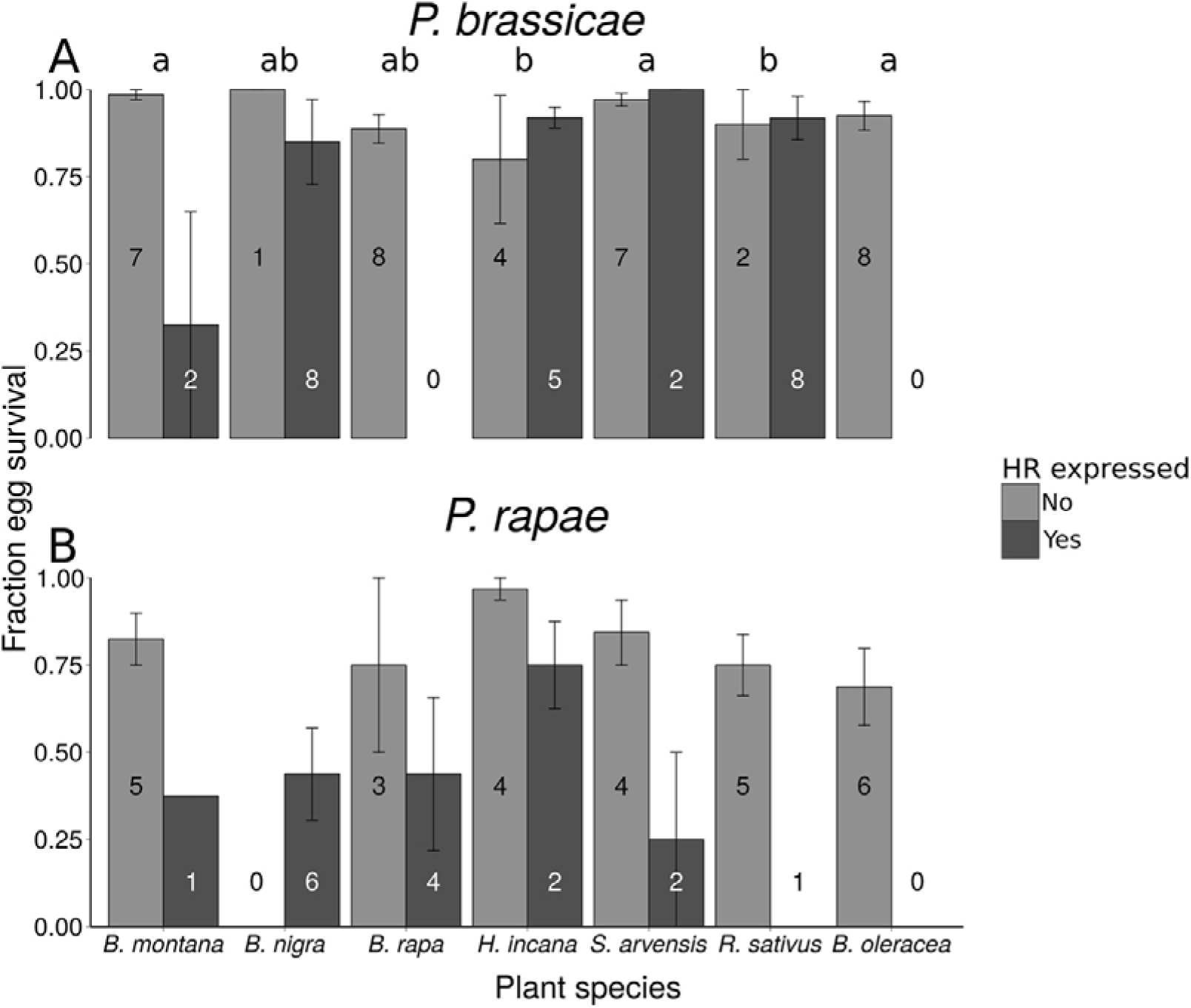
Effect of HR-like necrosis on survival of eggs of two *Pieris* species on different plant species (mean fraction ± SE). Numbers given in the bars indicate the number of plants. Different letters indicate significant differences (*P* <0.05, GLM) between plant species regardless of HR-like necrosis. (A) Fraction survival of *P. brassicae* eggs, with each egg clutch consisting of 20 eggs. (B) Fraction survival of *P. rapae* eggs, with eight eggs being laid per plant.

**Table 1:**
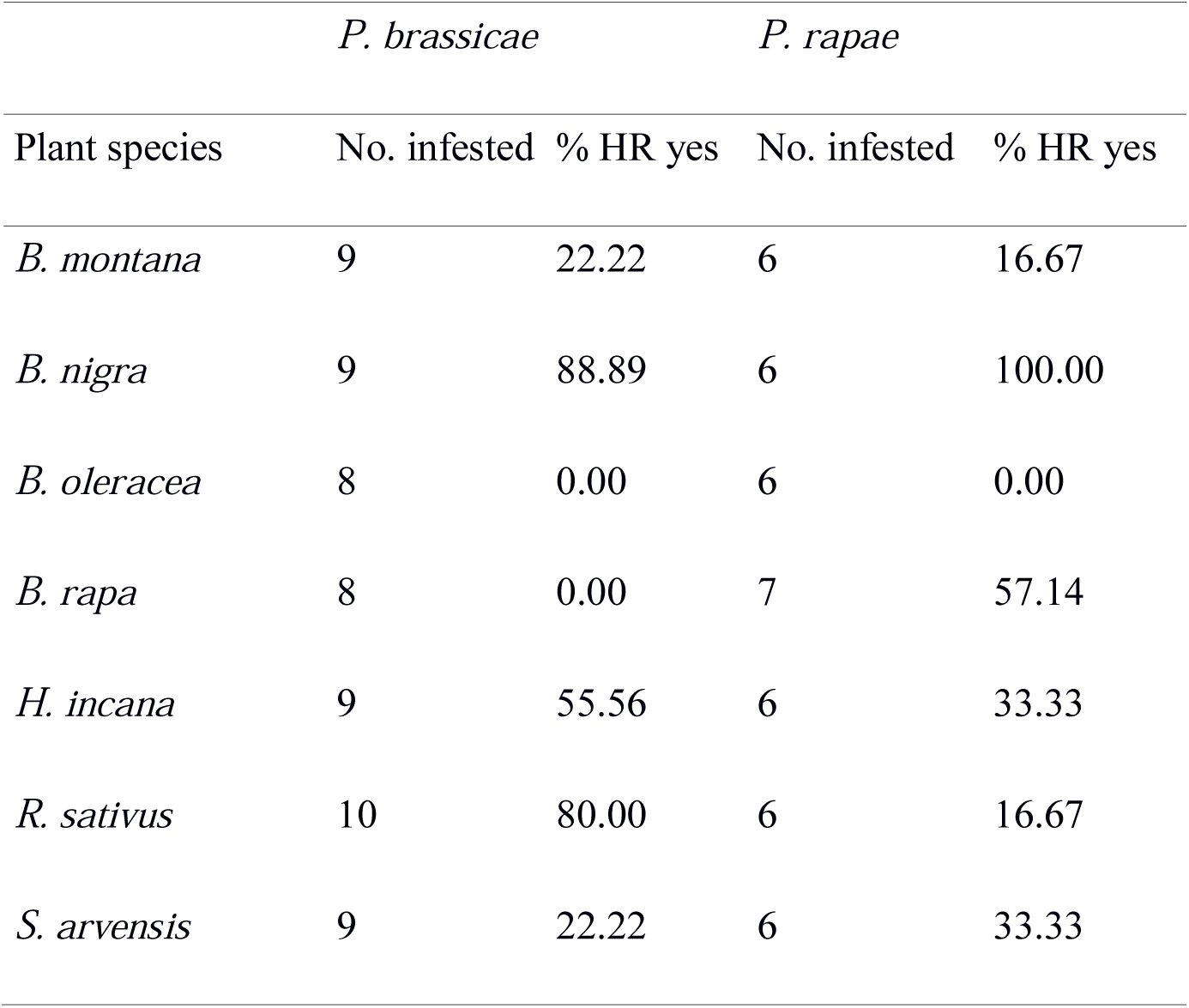
Number of infested plants and percentage of these plants expressing HR-like necrosis for both butterfly species separated for all plant species.

| Plant species | *P. brassicae* |  | *P. rapae* |  |
| --- | --- | --- | --- | --- |
|  | No. infested | % HR yes | No. infested | % HR yes |
| *B. montana* | 9 | 22.22 | 6 | 16.67 |
| *B. nigra* | 9 | 88.89 | 6 | 100.00 |
| *B. oleracea* | 8 | 0.00 | 6 | 0.00 |
| *B. rapa* | 8 | 0.00 | 7 | 57.14 |
| *H. incana* | 9 | 55.56 | 6 | 33.33 |
| *R. sativus* | 10 | 80.00 | 6 | 16.67 |
| *S. arvensis* | 9 | 22.22 | 6 | 33.33 |

#### Solitary species

The plant species selected by *P. rapae* females did not significantly affect egg survival (χ^2^ = 11.19, df = 6, *P* = 0.08, GLMM, Figure 3B). Six out of the seven tested plant species expressed HR-like necrosis induced by *P. rapae* eggs (Figure 3B, Table 1). A significantly higher fraction of *P. rapae* eggs survived on non-HR plants compared to plants expressing HR-like (χ^2^ = 13.58, df = 1, *P* < 0.001, GLMM, Figure 3B). This effect of *P. rapae* egg-induced HR-like necrosis on egg survival was – in contrast to the *P. brassicae* egg-induced response – independent of the plant species (χ^2^ = 4.43, df = 4, *P* = 0.35, GLMM).

### Correlation between oviposition preference and egg survival

When ignoring the expression of HR-like necrosis, we did not detect a significant correlation between oviposition preference and survival of the eggs for either of the two butterfly species (fraction of eggs laid related to egg survival; Spearman correlation; for *P. brassicae*: S = 32335, ρ = 0.10, *P* = 0.44, for *P. rapae*: S = 9086.3, ρ = 0.01, *P* = 0.97, Figure 4A). When looking at HR-like necrosis specifically, the fraction of *P. rapae* eggs laid was positively correlated with the fraction of plants expressing HR-like necrosis against those eggs (S = 4.57, ρ= 0.96, *P* < 0.001, Spearman correlation, see Figure 4B). For *P. brassicae*, this correlation was not significant (S = 67.72, ρ= 0.19, *P* = 0.64, Spearman correlation, see Figure 4B).

**Figure 4:**
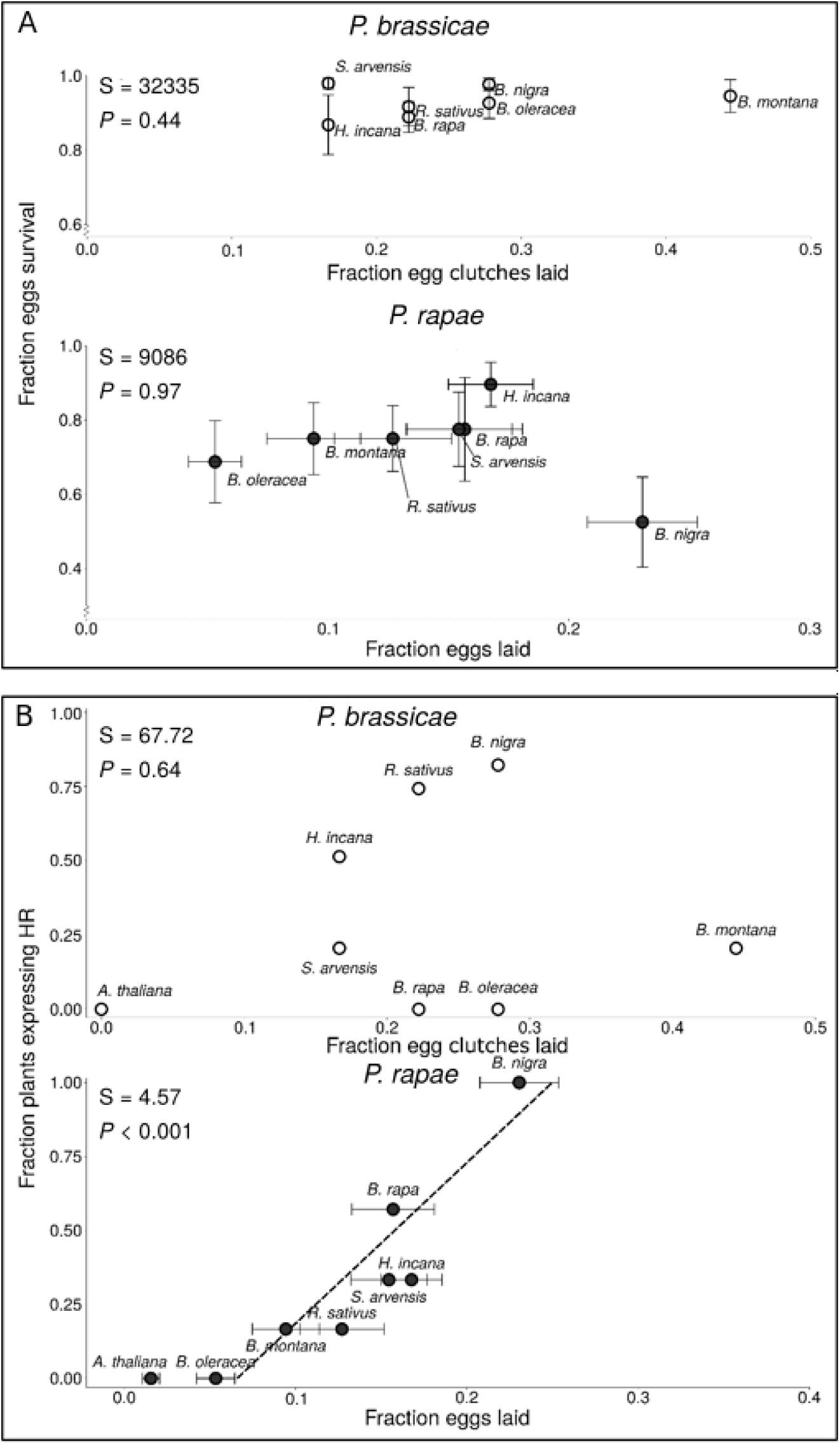
Correlation between fraction of eggs laid on different brassicaceous plants by two *Pieris* butterflies and (A) fraction of egg survival (mean ± SE) or (B) fraction of HR-like necrosis (mean ± SE). Text boxes show correlation results. Fraction of HR+ plants has no error bars, and for *P. brassicae*, no error bars for the preferences are available. N = 8-10 plants for *P. brassicae*, N = 5-6 plants for *P. rapae*.

### Effect of plant species, egg infestation and HR on larval performance

#### Gregarious species

The weight of seven-day-old *P. brassicae* caterpillars did not vary significantly depending on the plant species they were feeding on (χ^2^ = 12.44, df = 6, *P* = 0.05, LMM, Figure 5A). However, the plants’ response to prior egg deposition significantly affected performance of *P. brassicae* larvae. Seven-day-old larvae developing on plants that previously had received eggs (EF) performed significantly worse than those on plants that had not received eggs (F) (χ^2^ = 5.27, df = 1, *P* = 0.02, LMM, Figure 5B). This egg-mediated effect on anti-herbivore plant defense was independent of the plant species (no interactive effect between plant species and egg infestation on larval weight; χ^2^ = 2.51, df = 6, *P* = 0.87, LMM). HR-like necrosis induced by previously laid eggs did not affect the weight of caterpillars (χ^2^ = 0.72, df = 1, *P* = 0.40, LMM, see Figure 5C), and neither did plant species nor did the interaction between plant species and HR-like necrosis (χ^2^ = 5.76, df = 6, *P* = 0.45 and χ^2^ = 5.46, df = 4, *P* = 0.24, LMM).

**Figure 5:**
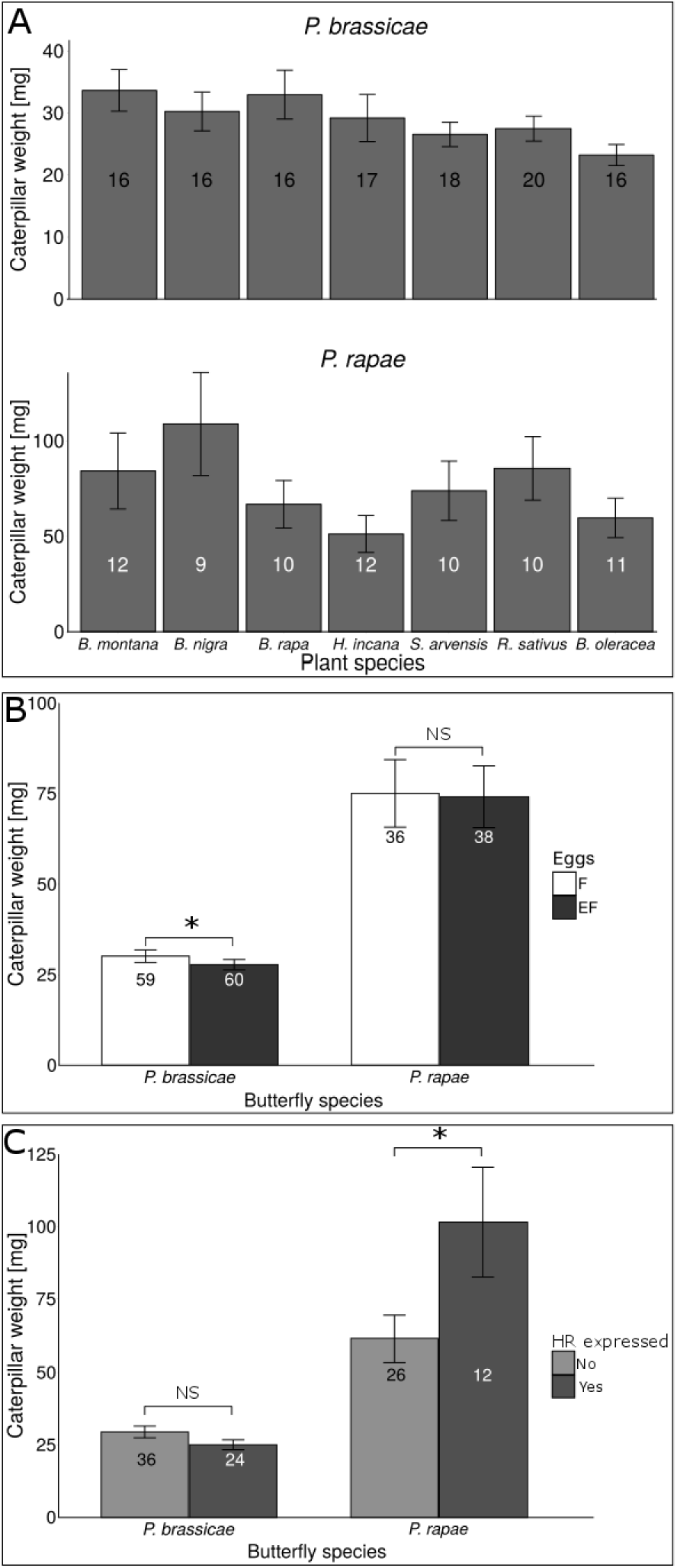
Effect of brassicaceous plant species (A), egg-mediated plant effects (B), and HR-like necrosis (C) on weight (mean ± SE) of seven-day-old *Pieris brassicae* or *P. rapae* caterpillars. In (A), weights of caterpillars feeding upon egg-free and previously egg-deposited plants are pooled. In (B), weights of caterpillars feeding upon egg-free and previously egg-deposited plants are shown separately. In (C), weights of caterpillars feeding upon previously egg-deposited plants are shown separately for plants expressing HR-like necrosis or not in response to egg deposition. The numbers in the bars represent the number of plants within the group. The weight of caterpillars was averaged per plant. Asterisks indicate significant differences. \**P* <0.05, ns: not significant, GLMM.

#### Solitary species

When considering seven-day-old *P. rapae* caterpillars on both egg-free and previously egg-deposited plants, their weight was not affected by the plant species they were feeding on (χ^2^ = 5.04, df = 6, *P* = 0.54; LMM, Figure 5A). When excluding the occurrence of HR-like necrosis induced by egg deposition, egg infestation preceding larval feeding did not affect larval weight (χ^2^ = 0.001, df = 1, *P* = 0.97; LMM, Figure 5B). Neither did the interaction between egg infestation and plant species affect larval weight (χ^2^ = 1.09, df = 6, *P* = 0.98, LMM, Figure 5B). Yet, larvae feeding on EF plants expressing an HR-like necrosis were significantly heavier than those feeding on EF plants that did not show HR-like necrosis (χ^2^ = 4.14, df = 1, *P* = 0.04, LMM, Figure 5C). Neither plant species nor the interaction between plant species and HR-like necrosis affected caterpillar weight on previously egg-infested plants (χ^2^ = 3.73, df = 6, *P* = 0.71 and χ^2^ = 3.93, df = 3, *P* = 0.27, LMM).

### Correlation between oviposition preference and larval performance

To assess whether there was a correlation between adult oviposition preference and larval performance, we first analyzed the relationship between the fraction of eggs laid and the weight of three or seven-day-old caterpillars feeding on previously oviposited EF plants and egg-free F plants for each plant species.

#### Gregarious species

Weight of seven-day-old *P. brassicae* larvae significantly and positively correlated with the number of eggs laid. Seven-day-old *P. brassicae* larvae were the heaviest on those plant species that received most egg clusters (S = 15964000, ρ = 0.17, *P* < 0.001, Spearman correlation, Figure 6A). The fraction of egg clusters laid did not correlate with the weight of caterpillars feeding on egg-free plants (S = 34, ρ= 0.39, *P* = 0.40, Spearman correlation, Supplementary Figure S2A).

**Figure 6:**
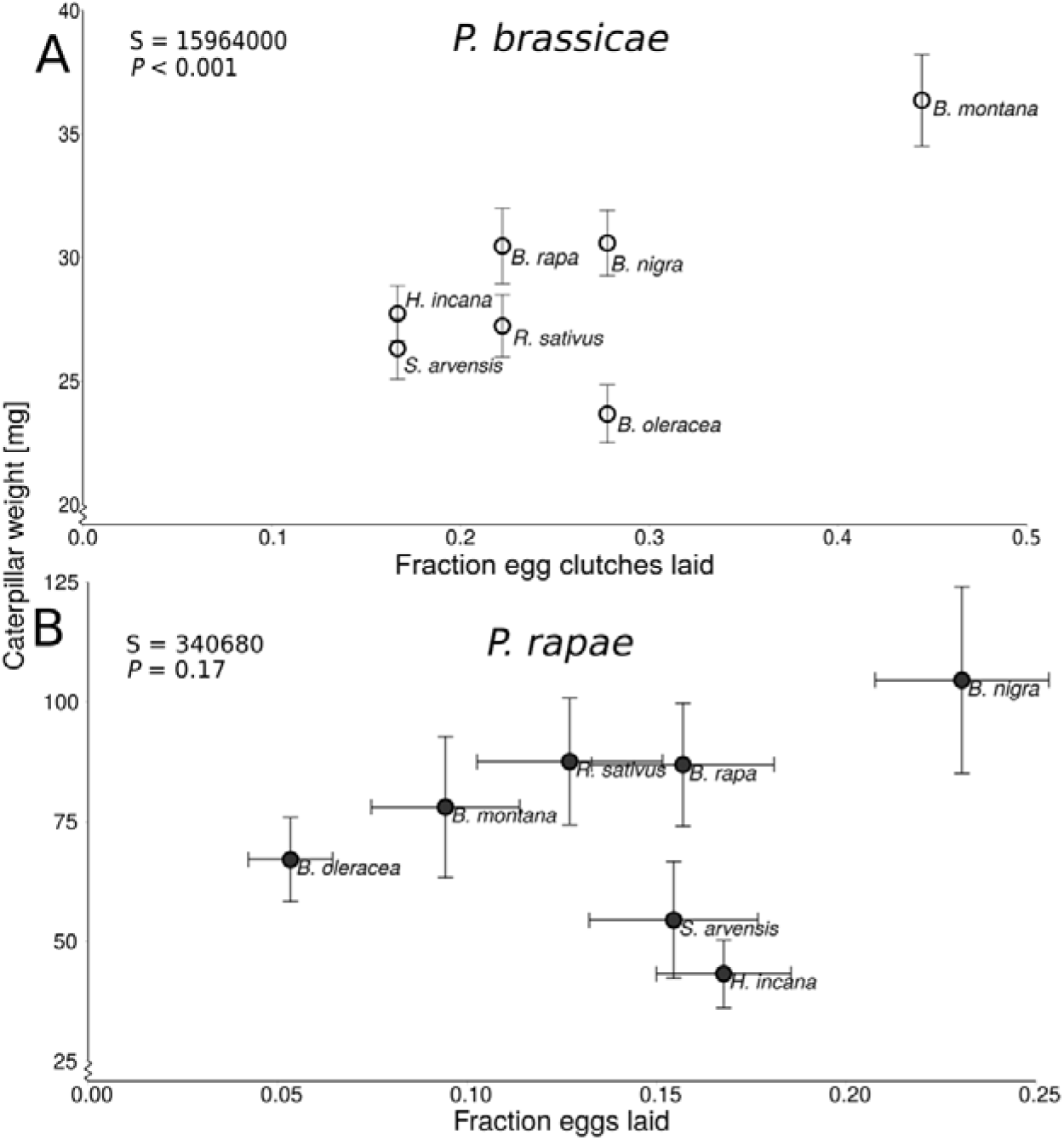
Correlation between oviposition preference and larval performance of seven-day-old *Pieris* caterpillars on different previously egg-infested EF brassicaceous plant species. Caterpillar weight (mean ± SE) and fraction of eggs/ egg clusters laid is shown. (A) Fraction of *Pieris brassicae* egg clutches laid per 18 test plants were used as preference measurement. (B) Fraction of *Pieris rapae* eggs laid per plant species (± SE) were used as preference data. Results of the Spearman correlation test are shown in text boxes. The y-axes do not start at zero to show the graph in greater detail.

When considering the weight of seven-day-old *P. brassicae* caterpillars with respect to the plant’s capability to express HR in response to the eggs, weight of caterpillars feeding on previously egg-deposited HR+ plants did not correlate with the fraction of egg clutches per plant (S = 22, ρ = −0.1, *P* = 0.95, Spearman correlation). Neither was a correlation found between the fraction of plants expressing HR-like necrosis in response to oviposition and the caterpillar weight (S = 36.65, ρ = 0.35, *P* = 0.45, Spearman correlation).

#### Solitary species

In contrast to *P. brassicae*, the weight of seven-day-old *P. rapae* larvae feeding on previously egg-infested plants did neither correlate with the fraction of eggs laid (S = 340680, ρ = 0.13, *P* = 0.17, Spearman correlation, Figure 6B) nor did larval weight correlate with egg load when larvae were feeding on egg-free plants (S = 50, ρ = 0.11, *P* = 0.84, Spearman correlation, Supplementary Figure S2B). Weight of seven-day-old caterpillars feeding on HR+ plants did not correlate with the fraction of eggs laid (S = 32, ρ = −0.6, *P* = 0.35, Spearman correlation). Furthermore, the weight of seven-day-old larvae was not correlated with the fraction of plants expressing HR-like necrosis (S = 29.52, ρ = 0.47, *P* = 0.28, Spearman correlation).

In addition to the weight of seven-day-old larvae, we also analyzed how the weight of three-day-old larvae is related to oviposition preference, to effects of prior egg deposition and expression of HR-like symptoms (compare Supplementary Figures S3, S4, and supplementary description of results). Two major differences were found when comparing the relationships of seven- and three-day-old larvae with the above-mentioned parameters. First, while weight of seven-day-old *P. brassicae* larvae and oviposition choices correlated significantly (Figure 6A), no such significant relationship was found when analyzing this relationship for three-day-old larvae and egg load (S = 30, ρ = 0.46, *P* = 0.30, Spearman correlation, Supplementary Figure S4A). As for seven-day-old caterpillars, the weight of three-day-old *P. rapae* larvae feeding upon HR-expressing plant species gained significantly more weight than larvae on non-HR-expressing plants (χ^2^ = 8.12, df = 1, *P* = 0.004, LMM, Supplementary Figure S3C). Plant species and interaction between both factors did not have any effect (χ^2^ = 7.44, df = 6, *P* = 0.28 and χ^2^ = 0.35, df = 3, *P* = 0.95, LMM, respectively). Correlation analysis for three-day-old *P. rapae* caterpillars did not reveal a significant correlation between oviposition choices and larval performances (S = 62, ρ = −0.11, *P* = 0.84, Spearman correlation, Supplementary Figure 4B).

## Discussion

The results of our study support the PPH when relating oviposition preferences and larval performances and show that oviposition-induced plant responses partly explain the PPH. Caterpillars of both species gained most biomass on those plant species that received most eggs. However, oviposition choices of both *Pieris* species do not correlate with egg survival. In response to singly laid *P. rapae* eggs, HR-like leaf necrosis was always induced in *B. nigra*, a plant species on which the eggs of this pierid species showed lowest survival rates. Unexpectedly, the solitary butterfly coped with this egg-induced plant defense by depositing most eggs on this plant species; *B. nigra* was most preferred for oviposition. Larval biomass of *P. rapae* was higher on plants expressing egg-induced HR-like necrosis compared to plants without necrosis. In contrast, the gregarious *P. brassicae* showed no significant oviposition preference for any of the tested plant species, and egg survival was hardly affected by HR-like necrosis. Weight of *P. brassicae* caterpillars feeding on previously egg-deposited plants was lower than of those feeding on egg-free plants. Such an egg-mediated plant effect on larval performance was not found in interactions with *P. rapae.* Hence, our data do not confirm the PPH when considering the relationship between oviposition preference and egg performances (egg survival). But we confirm the PPH for the relationship between preference and performance when considering the plant’s response to egg deposition that may affect larval performances.

Our finding that *P. rapae* females laid most eggs on a plant species (*B. nigra*) on which survival of eggs was lowest can hardly be considered an “oviposition mistake” (Larsson and Ekbom, 1995). Because *B. nigra* shows phenotypic variation in the expression of HR-like necrosis it might be difficult for the butterflies to discriminate between egg-resistant (HR+) and egg-susceptible (HR-) genotypes. Oviposition choices are influenced by different cues over long and short distances (Schoonhoven et al., 2005). Cues that signal intraspecific variation in suitability might be absent or of low detectability (Larsson and Ekbom, 1995). Yet, our data show that weight of *P. rapae* caterpillars was highest on *B. nigra* plants expressing HR where egg survival was reduced most. However, then the question arises why do the butterflies not lay most eggs on those plants where egg survival rates *and* larval performance are best? High egg survival rates on plants with a high egg load might result in several problems for the many hatching larvae, i.e. fast food depletion, easy detectability of caterpillars by parasitoids, increased cannibalism and the spread of pathogens (Prokopy and Roitberg, 2001). Therefore, the females might adjust their oviposition rate to the egg survival rate, which determines the extent of intraspecific competition, which caterpillars might experience on these plants.

Similar preferences and performances were obtained in studies with the polyphagous *Anastrepha ludens* fruit fly tested on six different host plants belonging to different families. The second most preferred plant species for oviposition, *Casimiroa edulis* (white sapote), was also the host on which larvae performed best. However, approximately half of all egg clutches laid on *C. edulis* were killed by a wound tissue growth response that led to egg encapsulation (Aluja et al., 2004; Birke and Aluja, 2018). We suggest that *P. rapae* butterflies can afford laying most eggs on the most nutritive plants, because here intraspecific competition among even numerous caterpillars is expected to be low due to the rich nutritional quality of the plant. While survival rates of *P. rapae* eggs on the most preferred host *B. nigra* were lowest, survival rates of *P. brassicae* eggs on the plant species with most egg depositions (*B. montana*) were similar to the plant species with fewer ovipositions. This suggests that *P. brassicae* does not adjust its oviposition choices to egg survival rates. *Pieris brassicae* might afford to be not choosy when selecting a host plant for oviposition because gregariously laid eggs have certain advantages with regard to egg survival over singly laid eggs. Gregariousness may for example contribute to protection from desiccation (Clark and Faeth, 1998; Stamp, 1980; Griese et al. 2017).

A positive relationship was found between the plant’s capability to respond to the singly laid *P. rapae* eggs by HR-like leaf necrosis and the oviposition preference of *P. rapae.* No such relationship was found between oviposition choices of *P. brassicae* and the occurrence of HR-like leaf necrosis in the various plant species. It is possible that those *P. rapae* eggs surviving potentially egg-killing plant responses harbored the fittest larvae that also face less competition, eventually leading to heavier growing larvae. Another possibility could be that caterpillars perform best on those plant species showing strong HR-like necrosis, because these plants provide high nutritional quality. Based on a meta-analysis, Wetzel et al. (2016) suggested that host plant nutritional quality might be more important for offspring performance than plant defenses against larvae; however, the analysis did not consider studies on plant defenses against insect eggs. Lastly, because less damage is inflicted to the leaf, less defenses might be induced, leading to heavier larvae.

Egg-mediated reinforcement of the plant’s defense against the caterpillars was only shown in the case of the gregarious *P. brassicae*, but not for the solitary *P. rapae.* When comparing weight of caterpillars on egg-free and previously egg-deposited plants (all species), *P. brassicae* caterpillars gained less weight on the latter. These results confirm previous results shown for *P. brassicae* (Bonnet et al., 2017; Geiselhardt et al., 2013; Pashalidou et al., 2015a; Pashalidou et al., 2015b; Pashalidou et al., 2015c; Pashalidou et al., 2013). Similarly, reinforced plant defense against insect larvae mediated by prior egg deposition has been shown in several interactions between plants and insects depositing eggs in clutches and feeding gregariously in early larval development, namely *Diprion pini* sawflies on pine (Beyaert et al., 2012); *Spodoptera littoralis* caterpillars on wild tobacco, (Bandoly et al., 2016; Bandoly et al., 2015) and *Xanthogaleruca luteola* leaf beetles on elm (Austel et al., 2016). Singly laid eggs of *P. rapae* did not prime plant defense against caterpillars. The reason why only plant responses to egg clutches of the gregarious *P. brassicae* negatively affected the performance of subsequently feeding caterpillars and not the singly laid eggs of *P. rapae* remains to be investigated.

Our study revealed that preferences and performances differ between the two butterfly species, which might be due to differences in oviposition modes of the butterfly species. The egg-laying mode might have affected egg survival when the plants expressed HR-like necrosis. When the plants received the singly laid eggs of *P. rapae*, we found a positive correlation between oviposition preference (fraction of eggs laid) and expression of HR-like necrosis (fraction of plants expressing HR). In contrast, no correlation was detected between these parameters when the plants received gregariously laid eggs of *P. brassicae.* In a previous study (Griese et al., 2017), we showed that expression of HR-like by *B. nigra* in response to *P. brassicae* did not affect survival of *P. brassicae* eggs when laid in clutches; however, when *P. brassicae* eggs were experimentally kept singly, they clearly suffered from low humidity at the oviposition site, which is characteristic of necrotic leaf tissue (Shapiro and DeVay, 1987). Hence, our current study supports the assumption that egg-induced HR-like has negative effects especially on survival of singly laid eggs (as those of *P. rapae*) rather than on clustered eggs (as those of *P. brassicae).* This is further supported by previous studies which showed that single eggs of *P. rapae* as well as of *P. napi* suffer high mortality when the host plant expresses HR-like necrosis (Fatouros et al., 2014; Shapiro and DeVay, 1987).

With respect to our initial questions, our study has shown that both *P. rapae* and *P. brassicae* laid most eggs on those plant species which provide best larval performance, while these plant species do not provide best egg survival rates. The plant species’ capability to activate HR-like necrosis in response to egg deposition affected the oviposition choice of *P. rapae* in so far, as this butterfly species laid most eggs on a plant species where HR-expression frequently occurred, and egg survival rates were low. This behavior might be considered a counter-adaptation because a high egg load on a plant with a high egg-killing capability ensures survival of at least some offspring. Since survival of eggs of *P. brassicae* was not affected by the plant’s HR-like, this butterfly species can afford laying many eggs on a plant species with high HR-expression frequency. Our data indicate that the gregarious oviposition mode of *P. brassicae* allows this butterfly species to be less choosy than *P. rapae* in selecting an oviposition site because the gregariously laid eggs are not affected by HR.

Future studies need to further address the question whether the differences in the effects of plant responses to these two pierid species are due to the different oviposition modes and larval feeding behaviors (singly vs. gregariously) or whether other insect species-specific traits are important as well. It remains to be investigated by studying interactions between more Brassicaceae and pierid species whether a gregarious oviposition mode as shown by *P. brassicae* and/or a high oviposition rate as shown by *P. rapae* laying single eggs on HR-expressing plant species may be considered as adaptations to the egg-induced leaf necrosis and whether these oviposition modes have evolved as countermeasures to this egg-inducible plant defense trait. Other countermeasures of pierid butterflies against egg-induced plant defenses include, e.g. oviposition on inflorescence stems instead of leaves, like observed in some other pierid species feeding on Brassicaceae (*P. napi* and *Anthocharis cardamines*) (NE Fatouros, personal observation) and feeding preference upon flowers if available (*A. cardamines* and *P. brassicae*) (Smallegange et al., 2007; Wiklund and Åhrberg, 1978). Because of the impact of insect oviposition on plant defense against hatching larvae shown in our and many other studies, we recommend to consider the egg phase when testing the PPH in herbivorous insects.

## Acknowledgements

We thank André Gidding, Léon Westerd, Joop Woelke and Frans van Aggelen for culturing insects and Unifarm of Wageningen University for providing plants and Rieta Gols for providing seeds used in the experiments.

This work was supported by The German Academic Exchange Service (DAAD) (57044990), the German Research Foundation (DFG, CRC 973; www.sfb973.de) and the Netherlands Organisation for Scientific Research (NWO) (NWO/ALW Veni grant no. 863.09.002 and NWO/TTW Vidi grant no. 14854).

## Author contribution statement

NEF, AP, FGP and EPI designed the experiments. EPI, AP and FGP performed the experiments. EG conducted statistical analysis and wrote the first draft of the paper. All authors interpreted results, drafted and revised the manuscript.

## Supporting information

Additional supporting information may be found online in the Supporting Information section at the end of the article.

**Results.** Effect of plant species, egg infestation and HR on performance of three-day-old larvae

**Table S1.** Results of the post hoc test on oviposition preference of *P. brassicae* and P. rapae differences between species.

**Table S2.** Results of post hoc test on egg survival of *P. brassicae* when compared between plant species.

**Table S3.** Results of single GLMs for each plant species tested for the influence of HR expression separated for *P. brassicae* on egg survival, used as a post hoc test.

**Figure S1.** Correlation between the number of eggs laid and plant fresh weight.

**Figure S2.** Correlation between egg laying preference and mea

**Figure S3.** Effect of brassicaceous plant species (A), egg-mediated plant effects (B), and HR-like necrosis (C) on weight (mean ± SE) of three-day-old *Pieris* caterpillars. n weight (± SE) of seven-day-old caterpillars on non-egg infested (F) plant species.

**Figure S4.** Linear regression between the fraction of eggs laid and the mass of 3-day old caterpillars (mean ± SE) on egg-infested (EF) plant species.

## References

Allen PE, 2010. Group size effects on survivorship and adult development in the gregarious larvae of *Euselasia chrysippe* (Lepidoptera, Riodinidae). Insect Soc 57:199–204.

Aluja M, Díaz-Fleischer F, Arredondo J, 2004. Nonhost status of commercial *Persea americana* ‘Hass’ to *Anastrepha ludens*, *Anastrepha obliqua*, *Anastrepha serpentina*, and *Anastrepha striata* (Diptera: Tephritidae) in Mexico. J Econ Entomol 97:293–309, 217.

Austel N, Eilers EJ, Meiners T, Hilker M, 2016. Elm leaves ‘warned’ by insect egg deposition reduce survival of hatching larvae by a shift in their quantitative leaf metabolite pattern. Plant Cell Environ 39:366–376.

Bandoly M, Grichnik R, Hilker M, Steppuhn A, 2016. Priming of anti-herbivore defense in *Nicotiana attenuata* by insect oviposition: herbivore-specific effects. Plant Cell Environ 39:848–859.

Bandoly M, Hilker M, Steppuhn A, 2015. Oviposition by *Spodoptera exigua* on *Nicotiana attenuata* primes induced plant defense against larval herbivory. Plant J 83:661–672.

Beyaert I, Köpke D, Stiller J, Hammerbacher A, Yoneya K, Schmidt A, Gershenzon J, Hilker M, 2012. Can insect egg deposition ‘warn’ a plant of future feeding damage by herbivorous larvae? P Roy Soc B-Biol Sci 279:101–108.

Birke A, Aluja M, 2018. Do mothers really know best? Complexities in testing the preference-performance hypothesis in polyphagous frugivorous fruit flies. B Entomol Res 108:674–684.

Bonnet C, Lassueur S, Ponzio C, Gols R, Dicke M, Reymond P, 2017. Combined biotic stresses trigger similar transcriptomic responses but contrasting resistance against a chewing herbivore in *Brassica nigra*. BMC Plant Biol 17:127.

Chew FS, Renwick JAA, 1995. Host plant choice in *Pieris* butterflies. Chemical Ecology of Insects 2 Boston, MA: Springer US. p. 214–238.

Clark BR, Faeth SH, 1997. The consequences of larval aggregation in the butterfly *Chlosyne lacinia*. Ecol Entomol 22:408–415.

Clark BR, Faeth SH, 1998. The evolution of egg clustering in butterflies: A test of the egg desiccation hypothesis. Evol Ecol 12:543–552.

Craig TP, Ohgushi T, 2002. Preference and performance are correlated in the spittlebug *Aphrophora pectoralis* on four species of willow. Ecol Entomol 27:529–540.

Denno R, Benrey B, 1997. Aggregation facilitates larval growth in the neotropical nymphalid butterfly *Chlosyne janais*. Ecol Entomol 22:133–141.

Desurmont GA, Weston PA, 2011. Aggregative oviposition of a phytophagous beetle overcomes egg-crushing plant defenses. Ecol Entomol 36:335–343.

Desurmont GA, Weston PA, Agrawal AA, 2014. Reduction of oviposition time and enhanced larval feeding: two potential benefits of aggregative oviposition for the viburnum leaf beetle. Ecol Entomol 39:125–132.

Edger PP, Heidel-Fischer HM, Bekaert M, Rota J, Glöckner G, Platts AE, Heckel DG, Der JP, Wafula EK, Tang M, Hofberger JA, Smithson A, Hall JC, Blanchette M, Bureau TE, Wright SI, de Pamphilis CW, Schranz ME, Barker MS, Conant GC, Wahlberg N, Vogel H, Pires JC, Wheat CW, 2015. The butterfly plant arms-race escalated by gene and genome duplications. PNAS 112:8362–8366.

Fatouros NE, Cusumano A, Danchin EGJ, Colazza S, 2016. Prospects of herbivore egg-killing plant defenses for sustainable crop protection. Ecol Evol 6:6906–6918.

Fatouros NE, Pineda A, Huigens ME, Broekgaarden C, Shimwela MM, Candia IAF, Verbaarschot P, Bukovinszky T, 2014. Synergistic effects of direct and indirect defenses on herbivore egg survival in a wild crucifer. Proc Roy Soc Lond B 281:20141254.

Feltwell DJ, 1982. Large white butterfly: The biology, biochemistry and physiology of *Pieris brassicae* (Linnaeus): Springer.

Fordyce JA, 2003. Aggregative feeding of pipevine swallowtail larvae enhances hostplant suitability. Oecologia 135:250–257.

Forister ML, 2004. Oviposition preference and larval performance within a diverging lineage of lycaenid butterflies. Ecol Entomol 29:264–272.

Forister ML, Nice CC, Fordyce JA, Gompert Z, 2009. Host range evolution is not driven by the optimization of larval performance: the case of *Lycaeides melissa* (Lepidoptera: Lycaenidae) and the colonization of alfalfa. Oecologia 160:551–561.

Friberg M, Posledovich D, Wiklund C, 2015. Decoupling of female host plant preference and offspring performance in relative specialist and generalist butterflies. Oecologia 178:1181–1192.

Geiselhardt S, Yoneya K, Blenn B, Drechsler N, Gershenzon J, Kunze R, Hilker M, 2013. Egg laying of cabbage white butterfly (*Pieris brassicae*) on *Arabidopsis thaliana* affects subsequent performance of the larvae. PLoS One 8:e59661.

Gols R, Bullock JM, Dicke M, Bukovinszky T, Harvey JA, 2011. Smelling the wood from the trees: Non-linear parasitoid responses to volatile attractants produced by wild and cultivated cabbage. J Chem Ecol 37:795.

Griese E, Dicke M, Hilker M, Fatouros NE, 2017. Plant response to butterfly eggs: inducibility, severity and success of egg-killing leaf necrosis depends on plant genotype and egg clustering. Sci Rep 7:7316.

Gripenberg S, Mayhew PJ, Parnell M, Roslin T, 2010. A meta-analysis of preference–performance relationships in phytophagous insects. Ecol Lett 13:383–393.

Harris MO, Sandanayaka M, Griffin W, 2001. Oviposition preferences of the Hessian fly and their consequences for the survival and reproductive potential of offspring. Ecol Entomol 26:473–486.

Hilker M, Fatouros NE, 2015. Plant responses to insect egg deposition. Annu Rev Entomol 60:493–515.

Hilker M, Fatouros NE, 2016. Resisting the onset of herbivore attack: plants perceive and respond to insect eggs. Curr Opin Plant Biol 32:9–16.

Hilker M, Schwachtje J, Baier M, Balazadeh S, Bäurle I, Geiselhardt S, Hincha DK, Kunze R, Mueller-Roeber B, Rillig MC, Rolff J, Romeis T, Schmülling T, Steppuhn A, van Dongen J, Whitcomb SJ, Wurst S, Zuther E, Kopka J, 2016. Priming and memory of stress responses in organisms lacking a nervous system. Biol Rev 91:1118–1133.

Hopkins RJ, Dam NMv, Loon JJAv, 2009. Role of glucosinolates in insect-plant relationships and multitrophic interactions. Annu Rev Entomol 54:57–83.

Jaenike J, 1990. Host specialization in phytophagous insects. Annu Rev Ecol Syst 21:243–273.

Karban R, 1983. Induced responses of cherry trees to periodical cicada oviposition. Oecologia 59:226–231.

König MAE, Wiklund C, Ehrlén J, 2016. Butterfly oviposition preference is not related to larval performance on a polyploid herb. Ecol Evol 6:2781–2789.

Larsson S, Ekbom B, 1995. Oviposition mistakes in herbivorous insects: Confusion or a step towards a new host plant? Oikos 72:155–160.

Lortzing V, Oberlander J, Lortzing T, Tohge T, Steppuhn A, Kunze R, Hilker M, 2018. Insect egg deposition renders plant defense against hatching larvae more effective in a salicylic acid-dependent manner. Plant Cell Environ 0:1–14.

Martínez G, Finozzi MV, Cantero G, Soler R, Dicke M, González A, 2017. Oviposition preference but not adult feeding preference matches with offspring performance in the bronze bug *Thaumastocoris peregrinus*. Entomol Exp Appl 163:101–111.

Mayhew PJ, 1997. Adaptive patterns of host-plant selection by phytophagous insects. Oikos 79:417–428.

Mayhew PJ, 2001. Herbivore host choice and optimal bad motherhood. Trends Ecol Evol 16:165–167.

Mazanec Z, 1985. Resistance of *Eucalyptus marginata* to *Perthida glyphopa* (Lepidoptera: Incurvariidae). Aust J Entomol 24:209–221.

Nufio CR, Papaj DR, 2001. Host marking behavior in phytophagous insects and parasitoids. Entomol Exp Appl 99:273–293.

Pashalidou FG, Fatouros NE, Van Loon JJA, Dicke M, Gols R, 2015a. Plant-mediated effects of butterfly egg deposition on subsequent caterpillar and pupal development, across different species of wild Brassicaceae. Ecol Entomol 40:444–450.

Pashalidou FG, Frago E, Griese E, Poelman EH, van Loon JJA, Dicke M, Fatouros NE, 2015b. Early herbivore alert matters: plant-mediated effects of egg deposition on higher trophic levels benefit plant fitness. Ecol Lett 18:927–936.

Pashalidou FG, Gols R, Berkhout BW, Weldegergis BT, van Loon JJA, Dicke M, Fatouros NE, 2015c. To be in time: egg deposition enhances plant-mediated detection of young caterpillars by parasitoids. Oecologia 177:477–486.

Pashalidou FG, Lucas-Barbosa D, van Loon JJA, Dicke M, Fatouros NE, 2013. Phenotypic plasticity of plant response to herbivore eggs: effects on resistance to caterpillars and plant development. Ecology 94:702–713.

Petzold-Maxwell J, Wong S, Arellano C, Gould F, 2011. Host plant direct defense against eggs of its specialist herbivore, *Heliothis subflexa*. Ecol Entomol 36:700–708.

Prokopy RJ, Roitberg BD, 2001. Joining and avoidance behavior in nonsocial insects. Annu Rev Entomol 46:631–665.

R Core Team, 2016. R: A language and environment for statistical computing. R Foundation for statistical computing. Vienna, Austria.

Reymond P, 2013. Perception, signaling and molecular basis of oviposition-mediated plant responses. Planta 238:247–258.

Root RB, Kareiva PM, 1984. The search for resources by cabbage butterflies (*Pieris rapae*): Ecological consequences and adaptive significance of markovian movements in a patchy environment. Ecology 65:147–165.

Scheirs J, Bruyn LD, Verhagen R, 2000. Optimization of adult performance determines host choice in a grass miner. Proc Roy Soc Lond B 267:2065–2069.

Schoonhoven LM, Van Loon JJA, Dicke M, 2005. Insect-Plant Biology. Oxford: Oxford University Press, UK.

Shapiro AM, DeVay JE, 1987. Hypersensitivity reaction of *Brassica nigra* L (Cruciferae) kills eggs of *Pieris* butterflies (Lepidoptera, Pieridae). Oecologia 71:631–632.

Sillén-Tullberg B, 1988. Evolution of gregariousness in aposematic butterfly larvae: A phylogenetic analysis. Evolution 42:293–305.

Smallegange RC, van Loon JJA, Blatt SE, Harvey JA, Agerbirk N, Dicke M, 2007. Flower vs. leaf feeding by *Pieris brassicae*: Glucosinolate-rich flower tissues are preferred and sustain higher growth rate. J Chem Ecol 33:1831–1844.

Stamp NE, 1980. Egg deposition patterns in butterflies: Why do some species cluster their eggs rather than deposit them singly? Am Nat 115:367–380.

Thompson JN, 1988a. Evolutionary ecology of the relationship between oviposition preference and performance of offspring in phytophagous insects. Entomol Exp Appl 47:3–14.

Thompson JN, 1988b. Variation in preference and specificity in monophagous and oliphagous swallowtail butterflies. Evolution 42:118–128. doi: 10.1111/j.1558-5646.1988.tb04112.x.

Thompson JN, Pellmyr O, 1991. Evolution of oviposition behavior and host preference in Lepidoptera. Annu Rev Entomol 36:65–89.

Wertheim B, Baalen E-JAv, Dicke M, Vet LEM, 2005. Pheromone-mediated aggregation in nonsocial arthropods: An evolutionary ecological perspective. Annu Rev Entomol 50:321–346.

Wetzel WC, Kharouba HM, Robinson M, Holyoak M, Karban R, 2016. Variability in plant nutrients reduces insect herbivore performance. Nature 539:425.

Wheat CW, Vogel H, Wittstock U, Braby MF, Underwood D, Mitchell-Olds T, 2007. The genetic basis of a plant–insect coevolutionary key innovation. PNAS 104:20427–20431.

Wiklund C, Åhrberg C, 1978. Host plants, nectar source plants, and habitat selection of males and females of *Anthocharis cardamines* (Lepidoptera). Oikos 31:169–183.

Wiklund C, Friberg M, 2008. Enemy-free space and habitat-specific host specialization in a butterfly. Oecologia 157:287–294.

Wiklund C, Friberg M, 2009. The evolutionary ecology of generalization: among-year variation in host plant use and offspring survival in a butterfly. Ecology 90:3406–3417.

