## Supplementary material for "Plant responses to butterfly oviposition partly explain preference-performance relationships on different brassicaceous species": Supplemtary data

### Results

*Effect of plant species, egg infestation and HR on performance of three-day-old larvae*

*Gregarious species*. Weight of three-day-old *P. brassicae* caterpillars varied in dependence of the plant species they were feeding upon ($\chi^2$ = 28.70, df = 6, $P$ <0.001, LMM, Figure S3A). However, the post hoc analysis did not show significant differences in weights of larvae feeding upon the various plant species. Neither did infestation of plants with eggs prior to larval feeding ($\chi^2$ = 0.03, df = 1, $P$ = 0.87, LMM) nor the interaction between the factors ′egg infestation′ and ′plant species′ ($\chi^2$ = 0.95, df = 6, $P$ = 0.99, LMM) affect weight of larvae three days after hatching (Figure S3B).

Furthermore, we tested whether HR-like necrosis induced by the egg deposition affects larval weight by considering only plants that previously received eggs (EF). No significant weight differences were found between caterpillars that fed on HR- or HR+ plants ($\chi^2$ = 0.04, df = 1, $P$ = 0.84, LMM, fig. 5C). Again, weights of larvae feeding on the tested plant species varied significantly ($\chi^2$ = 18.09, df = 6, $P$ = 0.006, LMM), but the linear hypothesis (post hoc) test did not show differences in weights of larvae feeding upon certain plant species. There was no significant interaction between the effects of ′plant species′ and ′HR′ on larval weight ($\chi^2$ = 5.28, df = 4, $P$ = 0.26, LMM).

*Solitary species*. Weight of three-day-old *P. rapae* caterpillars was neither affected by plant species they were feeding upon, nor by egg deposition prior to larval feeding nor by the interaction between plant species and egg infestation ($\chi^2$ = 5.31, df = 6, $P$ = 0.50; $\chi^2$ = 0.28, df = 1, $P$ = 0.60 and $\chi^2$ = 0.16, df = 6, $P$ = 1.00 respectively, LMM Figure S3A and B). However, the HR-like necrosis induced by egg deposition prior to larval feeding significantly affected larval performance. Larvae feeding on HR+ plants were significantly heavier than HR- plants ($\chi^2$ = 8.12, df = 1, $P$ = 0.004, LMM, Figure S3C). Plant species and interaction between plant species and HR- like expression did not affect caterpillar weight on plants previously exposed to eggs (EF) ($\chi^2$ = 7.44, df = 6, $P$ = 0.28 and $\chi^2$ = 0.35, df = 3, $P$ = 0.95, LMM).

**Table S1:** Results of the post hoc test on oviposition preference of *P. brassicae and P. rapae* differences between species.

| | Species comparison | z value | *P* |
|---|---|---|---|
| **P. brassicae** | *B. montana - A. thaliana* | 0.01 | 1.00 |
| | *B. nigra - A. thaliana* | 0.01 | 1.00 |
| | *B. oleracea - A. thaliana* | 0.01 | 1.00 |
| | *B. rapa - A. thaliana* | 0.01 | 1.00 |
| | *H. incana - A. thaliana* | 0.01 | 1.00 |
| | *R. sativus - A. thaliana* | 0.01 | 1.00 |
| | *S. arvensis - A. thaliana* | 0.01 | 1.00 |
| | *B. nigra - B. montana* | -1.03 | 0.96 |
| | *B. oleracea - B. montana* | -1.03 | 0.96 |
| | *B. rapa - B. montana* | -1.39 | 0.84 |
| | *H. incana - B. montana* | -1.75 | 0.61 |
| | *R. sativus - B. montana* | -1.39 | 0.84 |
| | *S. arvensis - B. montana* | -1.75 | 0.61 |
| | *B. oleracea - B. nigra* | 0.00 | 1.00 |
| | *B. rapa - B. nigra* | -0.38 | 1.00 |
| | *H. incana - B. nigra* | -0.80 | 0.99 |
| | *R. sativus - B. nigra* | -0.38 | 1.00 |
| | *S. arvensis - B. nigra* | -0.80 | 0.99 |
| | *B. rapa - B. oleracea* | -0.38 | 1.00 |
| | *H. incana - B. oleracea* | -0.80 | 0.99 |
| | *R. sativus - B. oleracea* | -0.38 | 1.00 |
| | *S. arvensis - B. oleracea* | -0.80 | 0.99 |

| | | |
|---|---|---|
| *H. incana - B. rapa* | -0.42 | 1.00 |
| *R. sativus - B. rapa* | 0.00 | 1.00 |
| *S. arvensis - B. rapa* | -0.42 | 1.00 |
| *R. sativus - H. incana* | 0.42 | 1.00 |
| *S. arvensis - H. incana* | 0.00 | 1.00 |
| *S. arvensis - R. sativus* | -0.42 | 1.00 |

| | **Species comparison** | **z value** | ***P*** |
|---|---|---|---|
| **P. rapae** | *B. montana - A. thaliana* | 6.67 | < 0.001 |
| | *B. nigra - A. thaliana* | 10.42 | < 0.001 |
| | *B. oleracea - A. thaliana* | 4.52 | < 0.001 |
| | *B. rapa - A. thaliana* | 8.48 | < 0.001 |
| | *H. incana - A. thaliana* | 8.87 | < 0.001 |
| | *R. sativus - A. thaliana* | 7.17 | < 0.001 |
| | *S. arvensis - A. thaliana* | 8.48 | < 0.001 |
| | *B. nigra - B. montana* | 7.19 | < 0.001 |
| | *B. oleracea - B. montana* | -3.08 | 0.04 |
| | *B. rapa - B. montana* | 3.22 | 0.03 |
| | *H. incana - B. montana* | 3.98 | 0.002 |
| | *R. sativus - B. montana* | 0.83 | 0.99 |
| | *S. arvensis - B. montana* | 3.22 | 0.02 |
| | *B. oleracea - B. nigra* | -9.42 | < 0.001 |
| | *B. rapa - B. nigra* | -4.26 | < 0.001 |
| | *H. incana - B. nigra* | -3.50 | 0.01 |
| | *R. sativus - B. nigra* | -6.49 | < 0.001 |
| | *S. arvensis - B. nigra* | -4.26 | < 0.001 |
| | *B. rapa - B. oleracea* | 6.02 | < 0.001 |
| | *H. incana - B. oleracea* | 6.69 | < 0.001 |

| | | |
|---|---|---|
| *R. sativus - B. oleracea* | 3.86 | 0.002 |
| *S. arvensis - B. oleracea* | 6.02 | < 0.001 |
| *H. incana - B. rapa* | 0.79 | 0.99 |
| *R. sativus - B. rapa* | -2.41 | 0.22 |
| *S. arvensis - B. rapa* | 0.00 | 1.00 |
| *R. sativus - H. incana* | -3.19 | 0.03 |
| *S. arvensis - H. incana* | -0.79 | 0.99 |
| *S. arvensis - R. sativus* | 2.41 | 0.22 |

**Table S2:** Results of post hoc test on egg survival of *P. brassicae* when compared between plant species.

| Species comparison | z value | *P* |
|---|---|---|
| *B. nigra - B. montana* | 0.002 | 1.00 |
| *B. oleracea - B. montana* | -1.11 | 0.90 |
| *B. rapa - B. montana* | -1.87 | 0.43 |
| *H. incana - B. montana* | -4.22 | < 0.001 |
| *R. sativus - B. montana* | -3.82 | 0.002 |
| *S. arvensis - B. montana* | -0.90 | 0.96 |
| *B. oleracea - B. nigra* | -0.002 | 1.00 |
| *B. rapa - B. nigra* | -0.002 | 1.00 |
| *H. incana - B. nigra* | -0.002 | 1.00 |
| *R. sativus - B. nigra* | -0.003 | 1.00 |
| *S. arvensis - B. nigra* | -0.002 | 1.00 |
| *B. rapa - B. oleracea* | -1.40 | 0.75 |
| *H. incana - B. oleracea* | -4.68 | < 0.001 |
| *R. sativus - B. oleracea* | -3.60 | 0.004 |
| *S. arvensis - B. oleracea* | -0.01 | 1.00 |
| *H. incana - B. rapa* | -3.69 | 0.003 |
| *R. sativus - B. rapa* | -3.08 | 0.02 |
| *S. arvensis - B. rapa* | 0.79 | 0.98 |
| *R. sativus - H. incana* | -1.09 | 0.91 |
| *S. arvensis - H. incana* | 3.52 | 0.01 |
| *S. arvensis - R. sativus* | 3.26 | 0.01 |

**Table S3:** Results of single GLMs for each plant species tested for the influence of HR expression separated for *P. brassicae* on egg survival, used as a post hoc test. The level of significance was 0.05. Dashes were used to indicate that the test was not applicable, because the plant species never expressed HR when exposed to *P. brassicae* eggs.

| | *P. brassicae* | | |
|---|---|---|---|
| Plant species | $\chi^2$ | df | *P* |
| *B. montana* | 19.50 | 1 | < 0.001 |
| *B. nigra* | 0.32 | 1 | 0.57 |
| *B. rapa* | --- | - | --- |
| *H. incana* | 0.70 | 1 | 0.40 |
| *S. arvensis* | 1.38 | 1 | 0.24 |
| *R. sativus* | 0.02 | 1 | 0.89 |
| *B. oleracea* | --- | - | --- |
| *A. thaliana* | --- | - | --- |

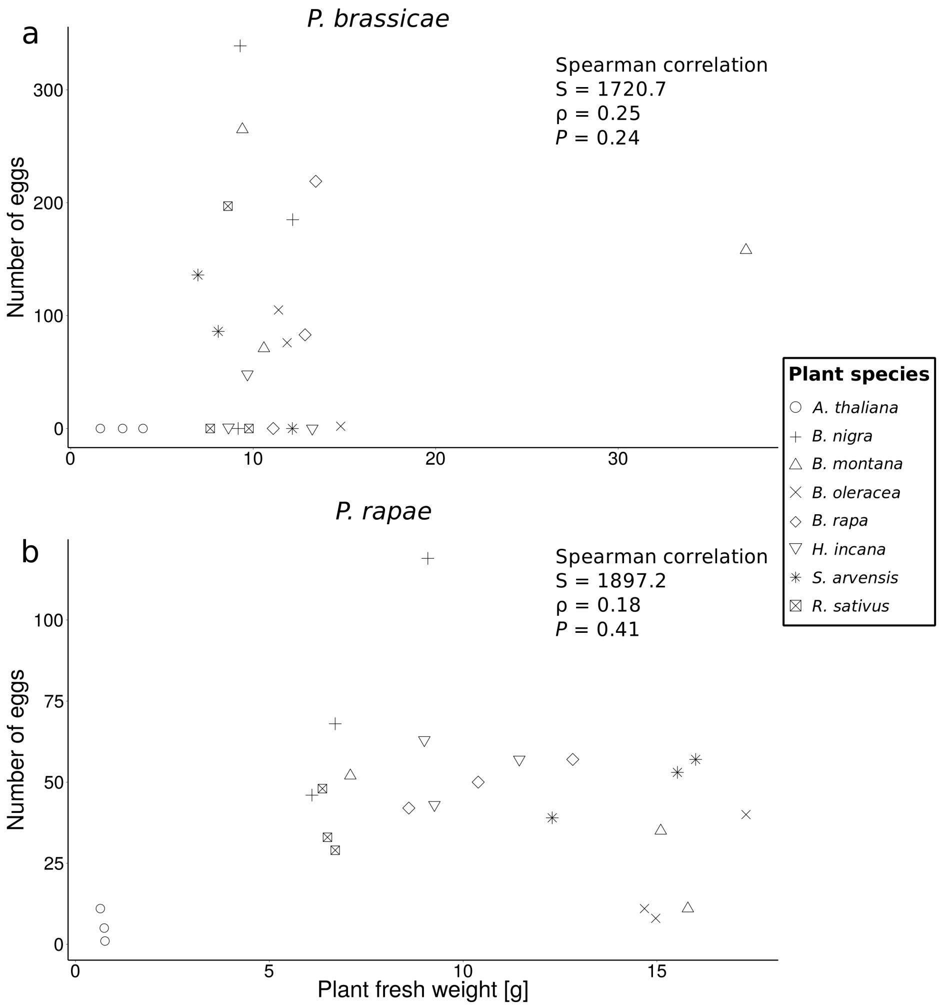


**Figure S1:** Correlation between the number of eggs laid and plant fresh weight. Neither for *P. brassicae* nor for *P. rapae* there is a significant correlation between the two factors.

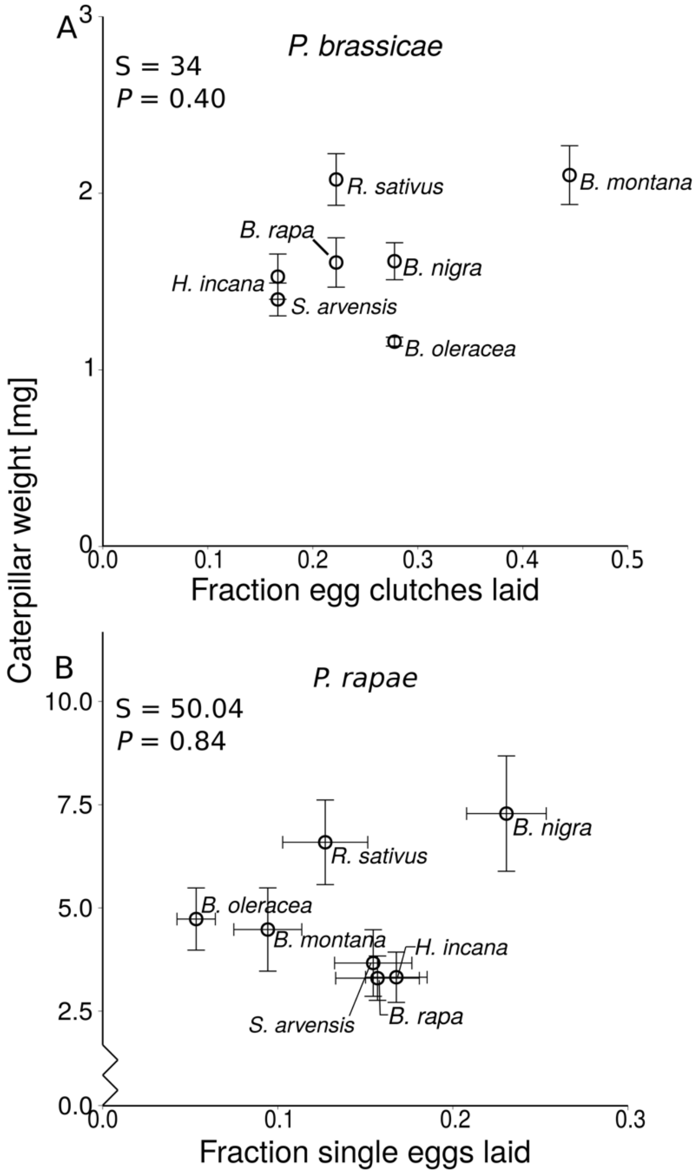


**Figure S2:** Correlation between egg laying preference and mean weight (± SE) of seven-day-old caterpillars on non-egg infested (F) plant species. A: *Pieris brassicae* correlation, the fraction of egg clutches laid per 18 test plants were used as preference measurement. B: *Pieris rapae* correlation mean eggs laid per plant species (± SE) were used as preference data. Neither correlation was significant (Spearman correlation).

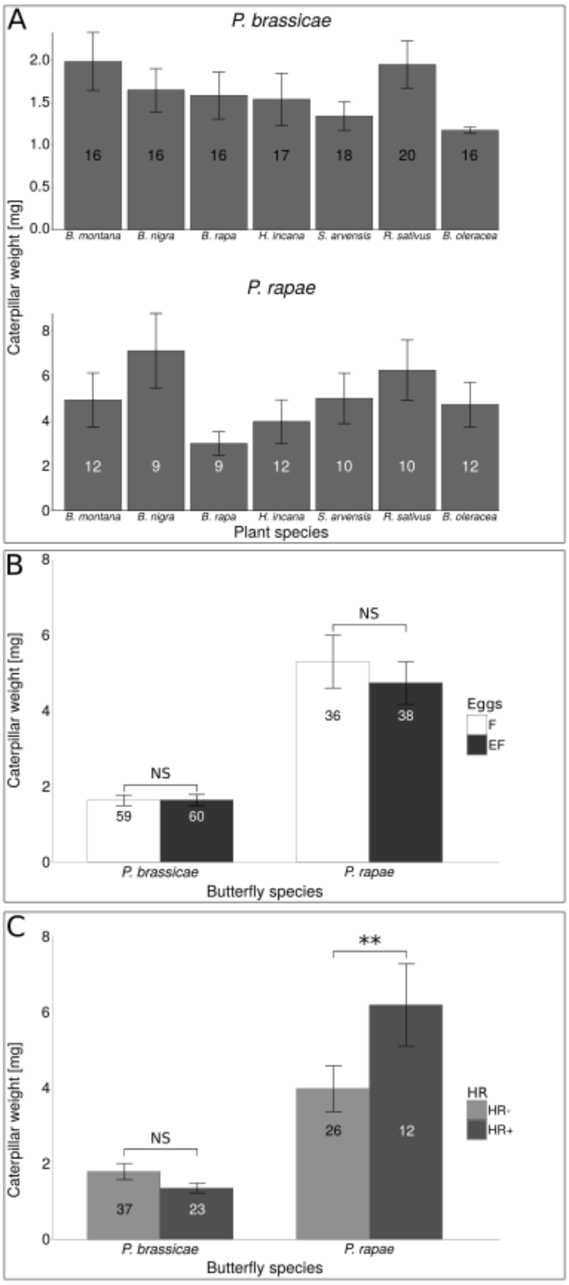


**Figure S3:** Effect of brassicaceous plant species (A), egg-mediated plant effects (B), and HR-like necrosis (C) on weight (mean ± SE) of three-day-old *Pieris* caterpillars. In (A), weights of caterpillars feeding upon egg-free and previously egg-deposited plants are pooled. In (B), weights of caterpillars feeding upon egg-free and previously egg-deposited plants are shown separately. In (C), weights of caterpillars feeding upon previously egg-deposited plants are shown separately for plants expressing HR-like necrosis or not in response to the egg deposition. The numbers in the bars represent the number of plants within the group. The weight of caterpillars was averaged per plant. Asterisks indicate significant differences. ***P* <0.01, ns: not significant, GLMM.

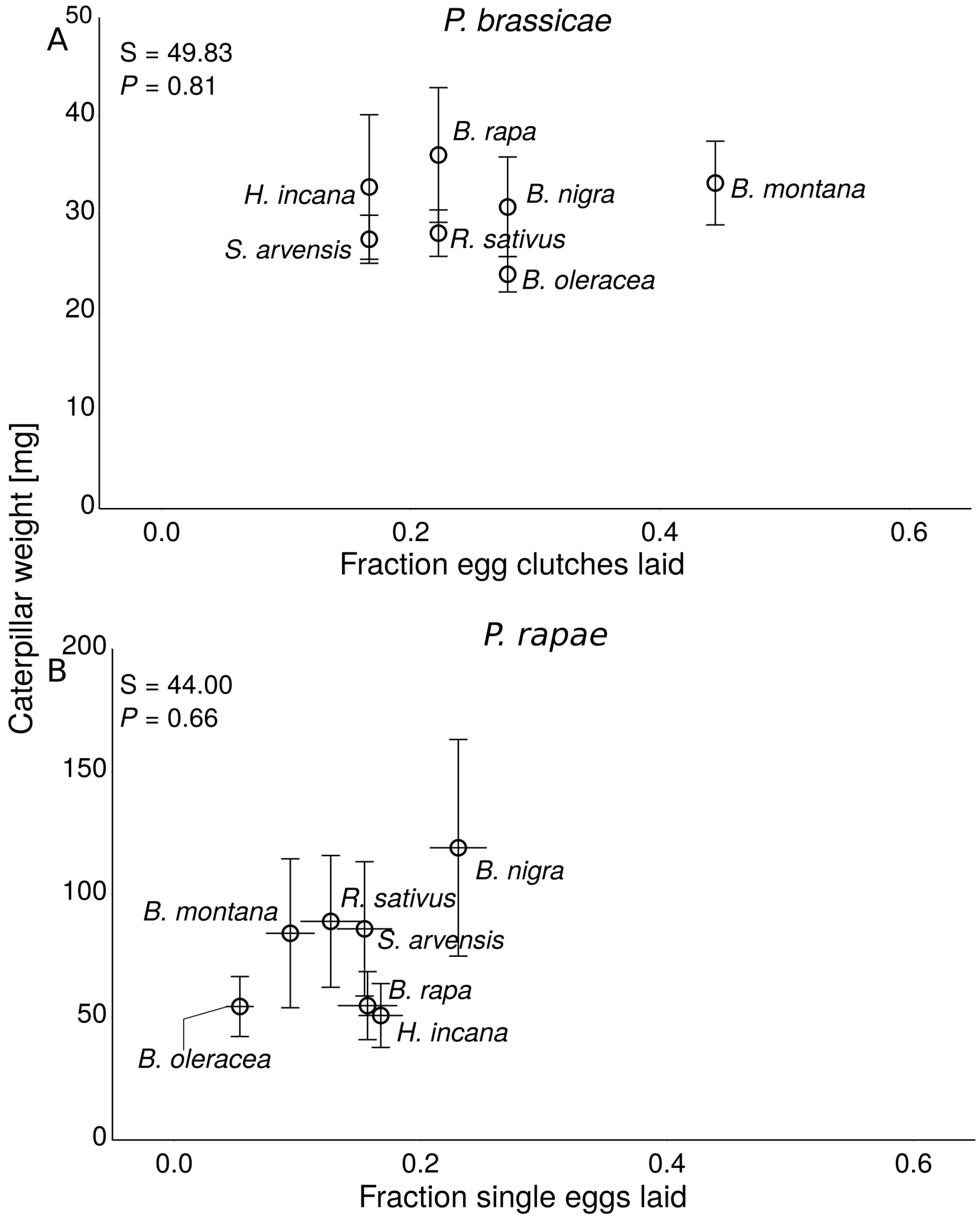


**Figure S4:** Linear regression between the fraction of eggs laid and the mass of 3-day old caterpillars (mean ± SE) on egg-infested (EF) plant species. A: *Pieris brassicae* correlation, the fraction of egg clutches laid per 18 test plants were used as preference measurement. B: *Pieris rapae* correlation mean eggs laid per plant species (± SE) were used as preference data.
